# Versatile patterns in the actin cortex of motile cells: Self-organized pulses can coexist with macropinocytic ring-shaped waves

**DOI:** 10.1101/2022.02.15.480577

**Authors:** Arik Yochelis, Sven Flemming, Carsten Beta

**Affiliations:** Department of Solar Energy and Environmental Physics, Blaustein Institutes for Desert Research, Ben-Gurion University of the Negev, Sede Boqer Campus, Midreshet Ben-Gurion 8499000, Israel; Department of Physics, Ben-Gurion University of the Negev, Be’er Sheva 8410501, Israel; Institute of Physics and Astronomy, University of Potsdam, Potsdam, 14476, Germany

## Abstract

Self-organized patterns in the actin cytoskeleton are essential for eukaryotic cellular life. They are the building blocks of many functional structures that often operate simultaneously to facilitate, for example, nutrient uptake and movement of cells. However, to identify how qualitatively distinct actin patterns can coexist remains a challenge. Here, we use bifurcation theory to reveal a generic mechanism of pattern coexistence, showing that different types of wave patterns can simultaneously emerge in the actin system. Our theoretical analysis is complemented by live-cell imaging experiments revealing that narrow, planar, and fast-moving excitable pulses may indeed coexist with ring-shaped macropinocytic actin waves in the cortex of motile amoeboid cells.

---

In biological cells, many functional processes take place simultaneously. Key examples are observed in the actin cytoskeleton of eukaryotic cells, which is a dense, dynamic biopolymer network located at the inner face of the plasma membrane. It provides the basis for important cellular functions, such as nutrient uptake, motility, and division. They rely on self-organized space-time patterns of cortical activity [1–6], both at the level of the actin cytoskeleton and in the upstream biochemical and mechanical signaling pathways [7– 16]. These patterns often appear simultaneously and are abundantly observed in many cell types, such as neurons, dendritic cells, and neutrophils [17–19]. Thus, understanding how cells can generate and robustly maintain qualitatively different coexisting cortical patterns is fundamental not only to cellular life and its pathologies, such as cancer [17, 20], but can also serve as a roadmap to the emerging field of synthetic biology, for which minimal candidate systems are required to robustly mimic intracellular spatiotemporal behaviors [21].

While recent research has primarily addressed the emergence of cortical actin waves that exhibit well defined wavelengths or temporal periods [18, 19], the coexistence of different patterns has been largely ignored and remains poorly understood. In this Letter, we use bifurcation theory to uncover a generic mechanism that relies on actin conservation, by which excitable pulses and oscillatory traveling waves emerge and coexist in the cell cortex. We then present experimental evidence from microscopy recordings of giant amoeboid cells, showing that different types of dynamical wave patterns can be indeed simultaneously maintained in the actin cortex of living cells.

## Actin conserved reaction-diffusion kinetics

Intracellular actin dynamics involves multiple spatiotemporal feedbacks ranging from the molecular level of protein kinetics to mechanical membrane deformation [18, 19, 22–25]. Yet, it has already been shown that fundamental insights into actin wave dynamics can be obtained by approximate descriptions in terms of reaction-diffusion (RD) systems that reduce the complexity of the actin cortex to a small number of key components and incorporate their most salient interactions [26]. These include a conserved pool of actin that is either encountered in its monomeric form (G-actin) or may polymerize to filamentous cortical structures (F-actin). In addition, F-actin may autocatalytically enhance its own formation, which reflects the well-known branching mechanisms and positive feedback to the activatory components of the upstream signaling pathway, such as PI3K and Ras [27]. On the other hand, the growth of filamentous structures is counterbalanced by inhibitory regulators, such as Coronin or Aip1 [28]. This is typically taken into account by an effective inhibitory component that is triggered in the presence of F-actin and downregulates the rate of polymerization. An overall schematic representation of the components and their interactions is displayed in Fig. 1.

**FIG. 1.**
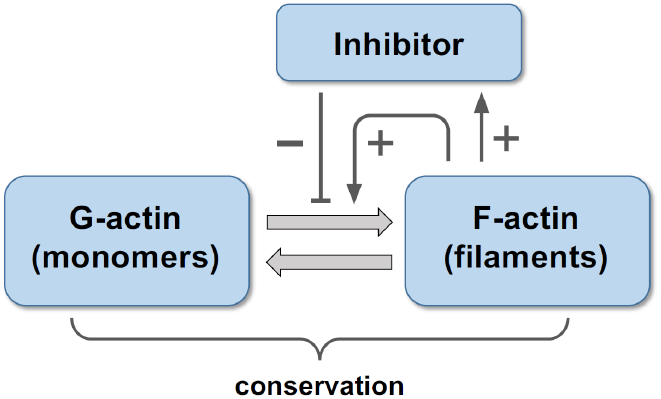
Schematic representation of interaction network of the three components of the actin dynamics model [29]. Mass conserved reactions are represented by thick light grey arrows. Activatory (+), and inhibitory (−) interactions are represented by thin dark grey arrows.

In general, due to autocatalytic growth and nonlinear feedback, multiple stable states may emerge that are characterized by different F-actin concentrations. From the study of circular dorsal ruffles (CDRs), it is known that at least two stable uniform states can coexist (bistability) [26]: a state that shows only a small amount of filamentous actin and is consequently dominated by a high G-actin concentration and a state of high F-actin concentration. In this case, also an unstable state at intermediate F-actin levels must exist but cannot be observed in experiments.

## Emergence of waves and pulses

To explicitly address the coexistence of traveling waves (TWs) and excitable pulses (EPs), we consider, without loss of generality, a reduced actin model [29] (for details see SM) that can be seen as a specific realization of the reaction scheme presented in Fig. 1. We denote the F-actin concentration as *N* (*x, t*), G-actin as *S*(*x, t*), and the inhibitor as *I*(*x, t*), so that the model phase space is spanned by **P** = (*N, S, I*), and the three uniform solutions are 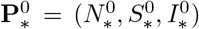 and 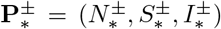 (see SM Eq. S3). In general, actin conservation implies that *L*^*−*1^∫_*L*_{*N* + *S*}d*x* = *A*, where *A* is the total actin concentration, and *L* is the domain length. In what follows, both will be used as control parameters. As G-actin monomers are inherently present, the G-actin-dominated state 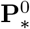 always exists and is linearly stable, while the two states 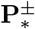 coexist for *A > A*_SN_ [29], where 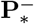 is linearly unstable, and 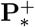 is linearly stable for *A > A*_W_; see the bifurcation diagram in Fig. 2(a,b). The condition *A > A*_W_ thus designates a bistable regime, where front solutions connecting 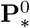 and 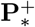 may form [26]. Here, we concentrate on the instabilities of the 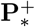 state that occur as *A* is decreased. The analysis is performed using the spatial dynamics methodology that combines bifurcation theory and numerical continuation [30], and the results are summarized in Fig. 2; see also SM for technical details.

**FIG. 2.**
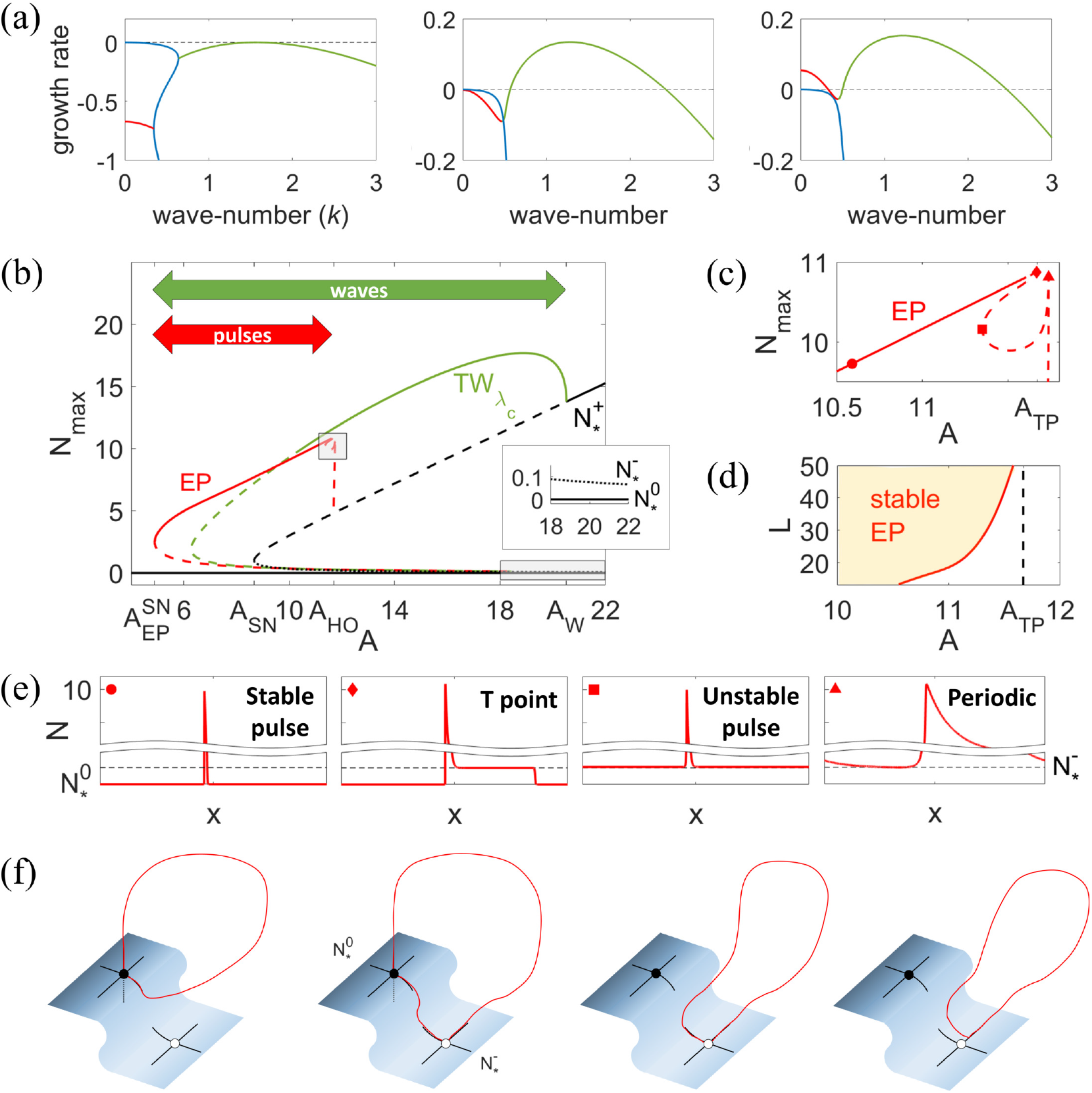
Analysis of following model system (S1) for waves and pulses (see also SM for details). (a) Perturbation growth rates at selected values of the total actin conservation, *A*. From left to right: the wave onset (green) at *A* = *A*_W_ ≃20.5, the wave unstable regime with homogeneous oscillations (HO) neutral (red) at *A*_HO_ ≃11.7, and an additional unstable regime at *A* = 11, where waves and HO are both unstable. The blue curve indicates real valued dispersion relations. (b) Bifurcation diagram showing the branches of primary traveling wave solutions 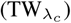 and excitable pulses (EPs) in periodic domains *L* = *λ*_*c*_ = 2*π/k*_*W*_ ≃ 4 and *L* = 1000, respectively. The branches are shown in terms of the maximal value of F-actin, *N* ; solid lines indicate the linear stability regions of the solutions along the branch with respect to multiple copies of the wavelength. The inset shows a zoom-in of the uniform solutions as indicated by the bottom shaded region. (c) Enlargement of the EP solutions about their origin near the T-point, *A* = *A*_TP_, (see top shaded region). (d) Stability of EPs with respect to the domain size, *L*. (e) EP profiles along the branch in (c), as indicated by respective symbols (f) Schematic (due to large scale separation) low-dimensional geometrical interpretation for EP profiles in (e).

As *A* is decreased, the uniform stable steady state 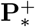 undergoes a wave (a.k.a. finite wavenumber Hopf) instability at *A*_W_, giving rise to traveling waves with wavenumber *k* = *k*_*W*_ ; see the dispersion relation in the left panel of Fig. 2(a). In the bifurcation diagram in Fig. 2(b), the primary 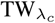 branch that emerges from *A*_W_ is displayed as a function of *A* (green line). Each point along the 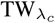 branch is associated with a periodic solution (with wavelength *λ*_*c*_ = 2*π/k*_*W*_) and plotted as the maximal value of the *N* component. These TW solutions bifurcate (super-critically) toward the unstable direction of 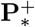 and are initially stable. As *A* is decreased below *A*_W_, additional TWs with larger spatial periods than *λ*_*c*_ emerge from the uniform state 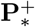, and their stability is also extended to lower *A* values; see SM Fig. S4. Similarly, standing waves (SWs) also bifurcate from *A*_W_, but we only mention their coexistence without explicitly showing their branch. Upon an increase of the domain size to cover multiple copies of TWs, a secondary instability of the Eckhaus-Benjamin-Feir type [31] destabilizes the TW solutions. For example, the instability onset for 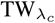 saturates at *A* ≃ 10.3 for *L* ∼ *O*(10*λ*_*c*_), similarly as reported in Yochelis *et al*. [32] (the instability point is marked in Fig. 2(b) by the transition from a solid to a dashed line). We note that this behavior is generic and not related to the specific choice of our model as it stems from a codimension-2 point, as discussed in more detail in SM.

An increase in the domain size is also related to the emergence of homogeneous oscillations (HO) at *A* = *A*_HO_ (in addition to TWs) for which the wave-number vanishes and the wavelength diverges; see the dispersion relation in the middle panel of Fig. 2(a). Numerically, it is impossible to implement infinite domains, but the typical approximation by periodic boundary conditions holds for such purposes. The continuation of periodic solutions from *A*_HO_, in a large domain *L* = 1000 ≫ *λ*_*c*_ (red line in Fig. 2(b)), shows that far from the onset, these solutions eventually develop into EPs. This evolution is depicted in Fig. 2(c), with the respective profiles along the branch in (c) displayed in panel (e). Up to the first sharp fold (homoclinic bifurcation in space) on the right (indicated by a triangle), the solutions are indeed periodic (with the period being as large as the domain size), and thus, at large amplitudes, they correspond to a broad pulse-like state; see Fig. 2(c) and the rightmost profile in (e). The branch then proceeds to the left and up to another fold (indicated by a rectangle), where the profile approaches a genuine pulse solution (see panel (e)), yet this solution is unstable since it is embedded in the background of the unstable 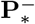 state as *x* → ±∞. Finally, after another fold at *A*_TP_ (indicated by a diamond), the branch extends to the left (toward 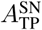), and within this range, stable EPs emerge (as shown, for example, at the location indicated by a dot). Notably, the latter are embedded in the background of 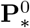, as shown by the respective leftmost profile in (e). The speeds of EPs are larger than the speeds of all TWs, as demonstrated in Fig. S5.

The transition from unstable to stable EP solutions corresponds to a generic global bifurcation that is also of codimension-2 and on an infinite line, also known as the T-point [33]. This bifurcation designates the merging of two EPs (homoclinic orbits in space), creating a bounded front state (heteroclinic cycle in space) connecting bidirectionally (biasymptotically) two uniform solutions 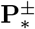 as *x* → ±∞ [34]. In Fig. 2(f), we schematically show the respective geometrical interpretation based on the profiles in (e). It is then straight-forward to see that while on large domains, the stability onset of EP is close to *A*_TP_, on small domains, where the excitation width of EP is on the order of the domain size, the onset shifts to lower *A* values, as shown in Fig. 2(d). This shift is associated with a structural change as the solutions depart from their solitary nature in space, becoming limit cycles (in space) rather than homoclinic orbits [33]; see also Figs. 2(e),(f). We refer the reader to the SM, specifically to Fig. S6, for further details on the pulse instability in domains of intermediate size, *O*(*λ*_*c*_) *< L < O*(10*λ*_*c*_), and the relation to pulse-splitting phenomena, cf. [29]. Next, we show that planar EPs can indeed coexist with other wave patterns in the cortex of amoeboid cells if their size is increased.

## Experimental observation of planar pulses

A well-established model organism to study the dynamics of the actin cortex is the social amoeba *Dictyostelium discoideum*. In *D. discoideum*, actin waves were reported more than 20 years ago [35], and they have been intensively studied with respect to their dynamics, structure, and biochemical composition [7, 8, 36, 37]. At the cytoskeletal level, these waves consist of traveling domains of increased F-actin concentration that are surrounded by a dense ring of F-actin. In terms of geometry and biochemical composition, they resemble planar macropinocytic patches, i.e. functional structures that facilitate cellular liquid uptake but cannot evolve into full three-dimensional cups due to the rigid cover slip surface [38, 39]. A particular advantage of the *D. discoideum* model system is the possibility to increase its cell size by electric-pulse-induced cell fusion, so that actin wave dynamics can be observed over large intact cell cortices independent of boundary effects [40]. In these giant cells, the macropinocytic waves take the form of actin-rich, band-shaped structures that travel at a characteristic speed and constantly split and coalesce in an irregular fashion [41–43].

These well-known wave patterns have been mostly described for axenic cell lines that feed on a liquid growth medium by macropinocytic liquid uptake. To explore new regimes of cytoskeletal dynamics, we changed the growth conditions and cultured the commonly used axenic *D. discoideum* lab strain AX2 together with bacteria. When feeding on bacteria, cells strongly enhance their pseudopod-driven motility in order to increase their chances of finding localized food sources [44]. Under these conditions, we expect pseudopod formation to compete with the formation of macropinocytic cups for the common actin pool. Also in this case, we observed macropinocytic actin waves that displayed the characteristic ring-shaped structure and meandering dynamics; see Fig. S8 and SM Movie 1. However, upon an increase in cell size, the cortical wave patterns exhibited an additional new feature that has not been observed in giant cells produced from axenically grown cells in the absence of bacteria. Namely, in addition to the well-known broad ring-shaped wave segments, the giant cells that were produced from cells grown on bacteria displayed sharp planar traveling pulses. These pulses behaved like genuine excitable solitary waves and have hitherto not been observed.

Both types of wave patterns — the broad macropinocytic traveling waves and the sharp planar pulses — coexisted in the same giant cell, and in Fig. 3(a), we show that they are distinct from each other in their profiles and propagation speeds, see also SM Movie 2. The planar pulses propagated at a higher speed than the broad traveling waves, and their profile was sharply peaked as compared to the wider profile of the traveling waves, which exhibited characteristic actin peaks at their leading and trailing edges. The latter results from the ring-shaped structure of the frustrated cup, see Fig. 3(b), a feature that is well known from earlier reports [41]. The planar pulses typically emerged near the cell border and traveled straight across the substrate-attached bottom membrane. Upon head-on collision, the planar pulses mutually annihilated as expected; see Fig. 3(c) and SM Movie 3 for an example.

**FIG. 3.**
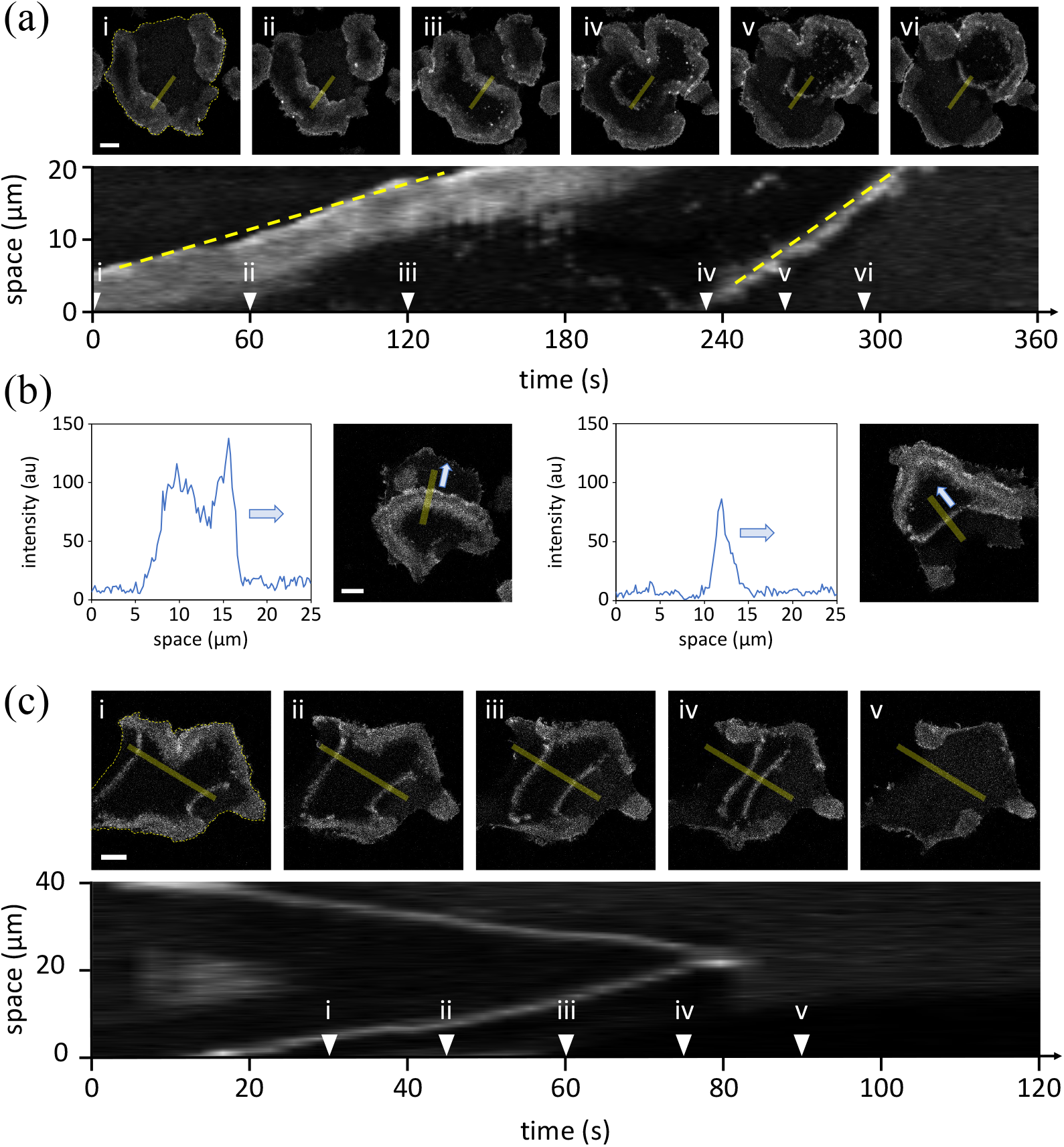
Coexisting waves and planar pulses in giant amoeboid cells. (a) Coexistence of slowly moving broad actin waves and rapidly propagating planar pulses in the cortex of a giant *D. discoideum* cell. The kymograph on the bottom row was taken along the yellow line displayed in the snapshots above. The dashed yellow lines are visual guides, indicating the different speeds of the wave and the pulse propagation. (b) Actin profiles of a wave (left) and a pulse (right) together with their respective snapshots; the yellow lines show the position where the profile was recorded, and the arrow indicates the direction of the wave and the pulse propagation, respectively. (c) Head-on collision and mutual annihilation of two counter-propagating planar pulses. The kymograph on the bottom row was taken along the yellow line displayed in the snapshots above. Dashed yellow lines in panels (i) of (a) and (c) are visual guides that indicate the cell outline. All scale bars correspond to 10 *µ*m.

## Discussion

Taken together, the rich and versatile spatiotemporal dynamics in the cell cortex is an essential part of actin-dependent cellular functions, such as phagocytosis, motility, and cell division. Nevertheless, the coexistence of different cortical patterns is poorly understood due to challenges in systematically deciphering the underlying mechanisms. For example, based on visual and/or statistical inspections alone, it is difficult to decide whether an observed spiral wave pattern results from an oscillatory, excitable, or bistable mechanism [18, 19].

In an attempt to address these fundamental mechanistic questions, we performed a bifurcation analysis of a mass conserved activator-inhibitor model of actin dynamics [26, 29], see Fig. 1, showing that even in a continuum-kinetic model that neglects most of the molecular details, essential features of pattern coexistence are revealed. Although we can envision other and more detailed models that may exhibit similar regimes of coexistence, the strength of our approach lies in uncovering the generic mechanism that leads to this behavior: a primary codimension-2 bifurcation and the evolution of homogeneous oscillations into excitable pulses via the so-called T-point (global) bifurcation in the presence of mass conservation. Specifically, using the total actin concentration as a control parameter, we revealed how recurrent traveling waves may coexist with planar excitable pulses; see Fig. 2. Following the theoretical analysis, we showed experimental recordings of giant *D. discoideum* cells, demonstrating that distinct actin wave patterns can indeed coexist in the cortex of living cells. Specifically, we found that, in addition to the well-known wide, ring-shaped macropinocytic waves, a novel type of narrow and planar, solitary pulse could be observed that revealed the typical excitable properties, see Fig. 3. In agreement with our theoretical predictions, these excitable pulses emerged upon an increase of the cell size and propagated faster than the ubiquitous traveling wave patterns. Besides the increased domain size, a lower total actin concentration is a key signature of this genuinely excitable regime.

We believe that, even though many more complex space-time patterns may coexist in cells [21], our results will provide a blueprint for analyzing and understanding fundamental questions of pattern coexistence based on bifurcation analysis and pattern formation theory. More broadly, our study of distinct coexisting wave types may also lead to a deeper understanding of cortical pathologies that could be related, for example, to cancerous phenotypes or uncontrolled tumor cell growth. It may furthermore inspire progress in the field of synthetic biology, where minimal, yet robust systems are required to reconstitute the essential features of self-organization in the cell cortex.

## Acknowledgments

We thank Kirsten Sachse for technical assistance and Alexander Shevchenko for helpful discussions. The research of CB has been partially funded by the Deutsche Forschungsgemeinschaft (DFG) – Project-ID 318763901 – SFB1294.

