## Supplementary Material for "Versatile patterns in the actin cortex of motile cells: Self-organized pulses can coexist with macropinocytic ring-shaped waves"

Arik Yochelis

*Department of Solar Energy and Environmental Physics,*

*Blaustein Institutes for Desert Research, Ben-Gurion University of the Negev,*

*Sede Boqer Campus, Midreshet Ben-Gurion 8499000, Israel and*

*Department of Physics, Ben-Gurion University of the Negev, Be'er Sheva 8410501, Israel*

Sven Flemming and Carsten Beta

*Institute of Physics and Astronomy, University of Potsdam, Potsdam, 14476, Germany*

(Dated: March 24, 2022)

### S1. MODEL EQUATIONS AND PARAMETERS

For the actin dynamics, we consider a three-component model (also see the text for details): $N(x,t)$ for F-actin, $S(x,t)$ for G-actin, and $I(x,t)$ for an inhibitor of actin polymerization. Following [1], the model equations in dimensionless forms read:

$$\frac{\partial N}{\partial t} = \frac{N^2 S}{1+I} - N + D_{\mathrm{N}}\frac{\partial^2 N}{\partial x^2}, \tag{S1a}$$

$$\frac{\partial S}{\partial t} = -\frac{N^2 S}{1+I} + N + D_{\mathrm{S}}\frac{\partial^2 S}{\partial x^2}, \tag{S1b}$$

$$\frac{\partial I}{\partial t} = r_{\mathrm{N}} N - r_{\mathrm{I}} I + D_{\mathrm{I}}\frac{\partial^2 I}{\partial x^2}. \tag{S1c}$$

To maintain fidelity with the experiments, we assume that $D_{\mathrm{I}} \ll D_{\mathrm{N}} < D_{\mathrm{S}}$ and $r_{\mathrm{I}} < r_{\mathrm{N}}$. For the analysis, we use the total number of actin monomers $A$ as a control parameter while keeping the rest of the parameters fixed (unless stated otherwise):

$$(D_{\mathrm{N}}, D_{\mathrm{S}}, D_{\mathrm{I}}, r_{\mathrm{N}}, r_{\mathrm{I}}) = (0.1, 1, 0.001, 2, 0.3)\,. \tag{S2}$$

### S2. LINEAR STABILITY OF UNIFORM AND NONUNIFORM SOLUTIONS

System S1 has three uniform solutions [1]; one is referred to as trivial due to the absence of the activator

$$\mathbf{P}_*^0 = (N_*^0, S_*^0, I_*^0) = (0, A, 0), \tag{S3a}$$

while the other two are finite

$$\mathbf{P}_*^\pm = (N_*^\pm, S_*^\pm, I_*^\pm) = (N_*^\pm, A - N_*^\pm, aN_*^\pm), \tag{S3b}$$

where $N_*^\pm = [A - a \pm \sqrt{(A-a)^2 - 4}]/2$ and $a = r_{\mathrm{N}}/r_{\mathrm{I}}$. While $\mathbf{P}_*^0$ always exists and, within the range of parameters considered here, is also stable, solutions $\mathbf{P}_*^\pm$ coexist for $A > A_{\mathrm{SN}} = a + 2$, where $\mathbf{P}_*^-$ is always unstable, and $\mathbf{P}_*^+$ is stable for $A > A_{\mathrm{W}}$. The linear stability onset of $\mathbf{P}_*^\pm$ is obtained following standard analysis to the linear order:

$$\mathbf{P}(x,t) - \mathbf{P}_*^\pm \propto e^{\sigma t + ikx}, \tag{S4}$$

where $\sigma$ is a growth rate of the respective wave-number $k$. By inserting (S4) into (S1) and expanding to the linear order, we obtain the dispersion relations in Fig. 2(a).

The existence of nonuniform solutions, such as traveling waves (TW) and excitable pulses (EP), was obtained using the numerical continuation package AUTO [2], after rewriting the system (S1) in a comoving coordinate frame $\xi = x + st$, where $s$ is the average translation speed at which these solutions propagate. The propagation speed is symmetric for left ($s > 0$) and right ($s < 0$) propagating solutions; hence, we employed periodic boundary conditions and only referred to one of these families. In

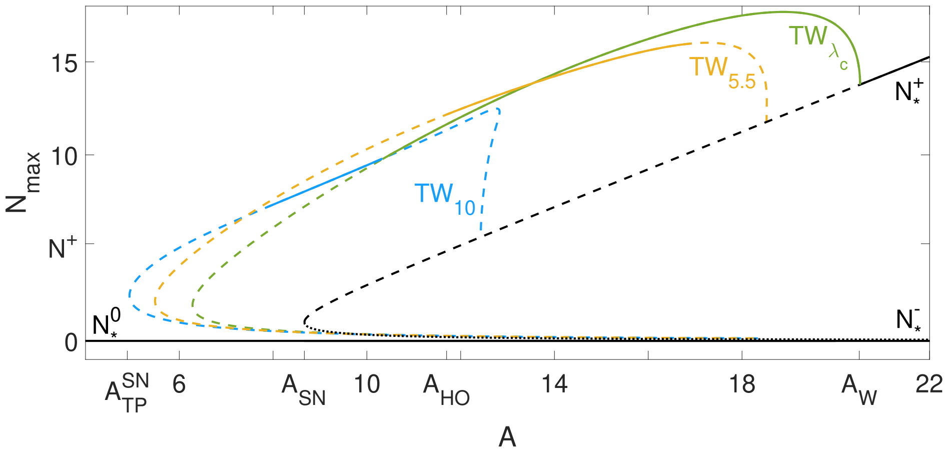


FIG. S1. Bifurcation diagram (as in Fig. 2(b)) showing two additional branches of traveling wave (TW) solutions with wavelengths $\lambda = 5.5$ (orange) and $\lambda = 10$ (cyan); solid lines indicate linear stability with respect to multiple copies of the wavelength, and the subscripts correspond to the wavelength. Parameters as in (S2).

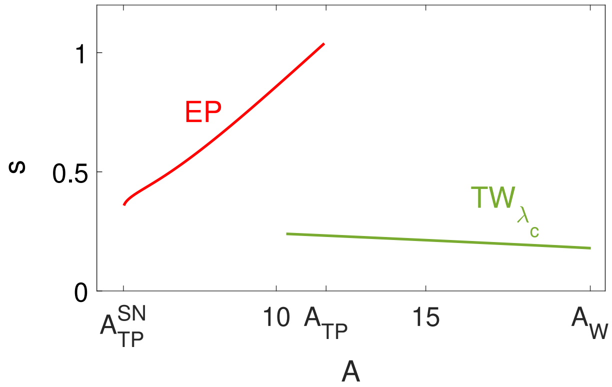


FIG. S2. Speed of stable EPs as compared to the speed of the critical TWs as a function of total actin concentration, as obtained numerically from continuation. Parameters as in Eq. S2.

this setting, TW and EP are steady state solutions, and to compute their stability, we retained the time derivatives employing a standard eigenvalue approach, with spatial derivatives being evaluated in the finite differences form. Linear stability

$$\mathbf{P}\left(\xi,t\right) - \mathbf{P}_{\mathrm{TW/EP}}\left(\xi\right) \propto e^{\sigma t}, \tag{S5}$$

is, however, determined for a large number of wavelengths (see also the main text), namely for $L = n\lambda$, where $\lambda$ is the TW wavelength, and $n$ is an integer, cf. [3]. In Figure S1, we show two representative TW solutions, $\lambda = 5.5$ (orange branch) and $\lambda = 10$ (cyan branch), together with the critical one denoted as TW$_{\lambda_c}$ (green branch). We note that $\lambda \to \infty$ as $A \to A_{\mathrm{HO}}$.

In an infinite domain, the stability of EPs persists from $A = A_{\mathrm{TP}} \approx A_{\mathrm{HO}}$ down to $A = A_{\mathrm{TP}}^{\mathrm{SN}}$ and their speed is larger than of TWs, as demonstrated in Fig. S2. Then the branch folds and continues as small amplitude unstable EPs. In practice, EP instability depends on the domain size (see Fig. 2(d)) and in the regime $O(\lambda_c) < L < O(10\lambda_c)$ it is triggered by the growth

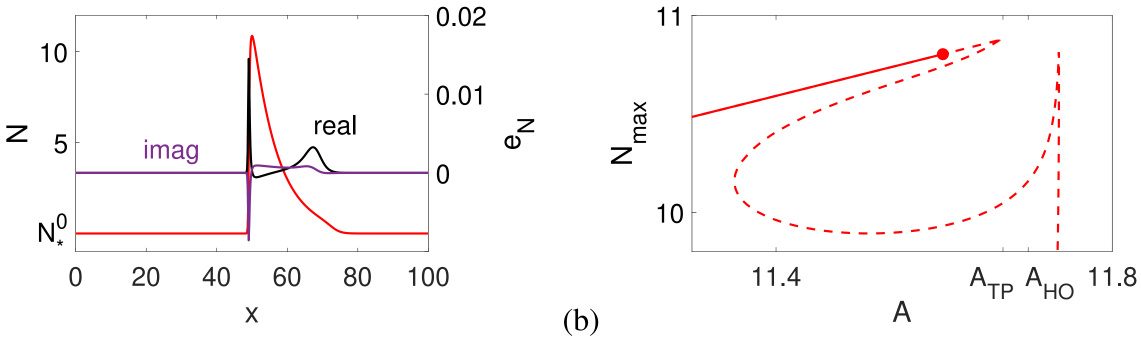


FIG. S3. Instability mechanism of excitable pulses. (a) Selected unstable quasi-excitable pulse (EP) profile, computed in domain $L = 100$ (red line), and the respective normalized eigenfunction ($e_{\rm N}$) computed at $A = 11.6$ near the T-point $A_{\rm TP}$, as shown by the bullet in (b). Parameters as in (S2).

of a shoulder at the rear of the pulse solution (due to the proximity to the T-point, as $A \to A_{\rm TP}$). We demonstrate this via the eigenfunction $e_{\rm N}$ of the first growing unstable eigenvalue. The real part of $e_{\rm N}$ exhibits a bumpy shape at the location of the pulse shoulder for $A \lesssim A_{\rm TP}$ (see Fig. S3). In the context of the collisions discussed in [1], the translational invariance is lost, and thus, since the perturbation in this range is of $O(1)$, the instability leads to the formation of standing waves that may resemble, but should not be confused with, the backfiring phenomenon [4–8].

### S3. AMPLITUDE EQUATIONS FOR THE UNIFORM OSCILLATIONS AND WAVE INSTABILITY WITH MASS CONSERVATION

For a multi-component system as described in Fig. 1, this occurs, most generally, via the oscillatory-wave codimension-2 bifurcation that requires the tuning of two parameters, here $A \to A_{\rm HO}$ and $D_{\rm N} \to D_{\rm W}$, as shown in Fig. S4. In this case, it is coupled to a large-scale mode (LSM), also known as the Goldstone mode, due to mass conservation. Weakly nonlinear theory (center manifold reduction) demonstrates that the patterns forming near such a codimension-2 point can be described by the following leading order approximation:

$$\mathbf{P} \sim \Big\{ \underbrace{B_0(X,T)e^{i\omega_0 t}}_{\text{HO}} + \underbrace{B_L(X,T)e^{i\omega_W t + ik_W x} + B_R(X,T)e^{i\omega_W t - ik_W x}}_{\text{waves}} + c.c. \Big\} + \underbrace{C(X,T)}_{\text{LSM}} + \text{h.o.t.}, \tag{S6}$$

where, $X, T$ are slow space and time scales as compared to $x, t$, respectively, so that complex amplitudes $B_0, B_{L/R}$ describe the weak spatiotemporal modulations of homogeneous oscillations (HO) associated with frequency $\omega = \omega_0$ and wavenumber $k = 0$, and the left/right propagating waves associated with $\omega = \omega_W$ and $k = k_W$ at the instability onset, respectively. The real valued field $C$ is associated with mass conservation, $c.c.$ stands for complex conjugated, and $h.o.t.$ stands for high order terms. In Fig. S4, the red branch represents the onset of homogeneous oscillations at $k = 0$, the green branch corresponds to the traveling wave instability at $k = k_W$, and the large-scale mode is represented in blue.

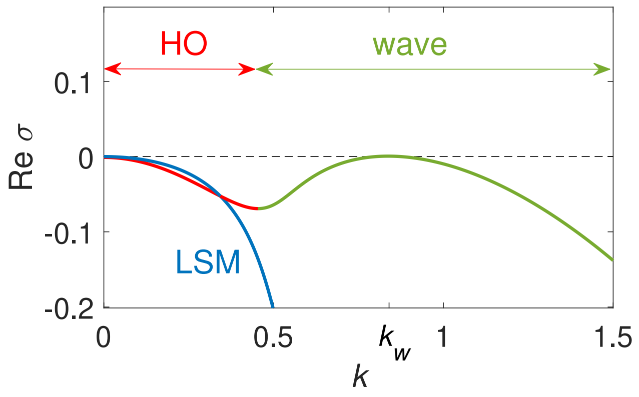


FIG. S4. Three growth rates associated with the real parts of the dispersion relations computed by the linearization of the mass conserved activator-inhibitor model (S1) about $\mathbf{P}_*^+$ at the codimension-2 point, $D_{\rm N} = D_{\rm W} \simeq 0.32$ and $A = A_{\rm HO} \simeq 11.7$ (other parameters as in (S2)): simultaneous onset of homogeneous oscillations (HO, red) and traveling waves (wave, green) in the presence of a large-scale mode (LSM, blue) that is associated with mass conservation, displaying a persistent neutral mode (at zero wave-number).

We note that partial representations along the lines of (S6) have been introduced without mass conservation in [9], while waves coupled to a conserved quantity (LSM) were initially introduced in [10]. The approximation (S6), however, is more general, exhibiting a seven-dimensional center manifold. Using standard symmetry considerations, the corresponding amplitude equations for $B_0$, $B_L$, $B_R$, and $C$ can also be specified in the weakly nonlinear regime

$$\partial_t B_0 = \overbrace{r_0 B_0 - \beta \left|B_0\right|^2 B_0 + \delta_0 \partial_x^2 B_0}^{\text{CGLE for HO}} - \overbrace{\gamma_L \left|B_L\right|^2 B_0 - \gamma_R \left|B_R\right|^2 B_0 - \gamma_C C B_0}^{\text{coupling}} - \overbrace{\mu B_L B_R \overline{B}_0}^{\text{resonance}}, \tag{S7a}$$

$$\left(\partial_t + s\partial_x\right) B_L = \overbrace{r_w B_L - b \left|B_L\right|^2 B_L - g_w \left|B_R\right|^2 B_L + \delta_w \partial_x^2 B_L}^{\text{CGLE for left TW}} - \overbrace{g_0 \left|B_0\right|^2 B_L - g_C C B_L}^{\text{coupling}} - \overbrace{m B_0^2 \overline{B}_R}^{\text{resonance}}, \tag{S7b}$$

$$\left(\partial_t - s\partial_x\right) B_R = \overbrace{r_w B_R - b \left|B_R\right|^2 B_R - g_w \left|B_L\right|^2 B_R + \delta_w \partial_x^2 B_R}^{\text{CGLE for right TW}} - \overbrace{g_0 \left|B_0\right|^2 B_R - g_C C B_R}^{\text{coupling}} - \overbrace{m B_0^2 \overline{B}_L}^{\text{resonance}}, \tag{S7c}$$

$$\partial_t C = \overbrace{\partial_x^2 \left[\delta_C C + \sigma_0 \left|B_0\right|^2 + \sigma_w \left(\left|B_L\right|^2 + \left|B_R\right|^2\right)\right]}^{\text{LSM}}, \tag{S7d}$$

where $r_0$ and $r_w$ are related to the distance from the HO and wave instability onsets, with the critical wavenumbers and frequencies as $(k, \omega) = (0, \omega_0)$ and $(k, \omega) = (k_W, \omega_k)$, respectively, $\delta_C, \sigma_0, \sigma_w$ are the real parameters, and all the others are complex. Each of the oscillatory amplitudes ($B_0$, $B_{L/R}$) in (S7) comprises the typical form of a complex Ginzburg-Landau equation (CGLE), a coupling term, and a term that may appear due to temporal resonances of $\omega_0$ and $\omega_k$. The parameters in (S7) depend on the parameters of the original system and will be discussed elsewhere.

### S4. CELL STRAINS, CULTURE CONDITIONS, AND TRANSFORMATION

The axenic AX2 strain of *D. discoideum* was cultivated on a bacterial lawn of *Klebsiella aerogenes* on SM agar plates (peptone 10 g/l, yeast extract 1 g/l, glucose 10 g/l, $KH_2PO_4$ 1.9 g/l, $K_2HPO_4 \cdot 3H_2O$ 1.3 g/l, $MgSO_4$ anhydrous 0.49 g/l, 1.7% agar) or in 10-cm dishes with Sörensen's buffer (8 g $KH_2PO_4$, 1.16 g $Na_2HPO_4$, pH 6.0) supplemented with 50 $\mu$M $MgCl_2$, 50 $\mu$M $CaCl_2$ and containing *K. aerogenes* at an $OD_{600}$ of 2 at 22°C as described by Paschke *et al.* [11]. The AX2 strain was adapted to growth in HL5 medium via macropinocytosis but it was still able to switch to growth based on phagocytosis of bacteria. To adapt them to these different growth conditions, the AX2 cells were cultivated for at least three days in a bacterial suspension or on SM agar plates with a bacterial lawn before they were used for the fusion protocol described below. A plasmid encoding for a Lifeact-GFP fusion protein was transformed into the AX2 cells by electroporation as previously described [12] with an ECM2001 electroporator (Harvard Apparatus, Holliston, MA, USA 01746-1388) using three square wave pulses of 500 V for 30 ms in electroporation cuvettes with a gap of 1 mm. Two plasmids for the expression of Lifeact-GFP were used: pDM138, which mediates G418 resistance [11], and pSF99, which mediates hygromycin resistance [12]. Transformants were selected with either 33 µg/ml hygromycin (Invivogen, France) or $\mu$g/ml G418 (VWR, Germany) in 10-cm dishes and usually appeared 2-4 days post-transformation.

### S5. CELL FUSION

Fusion of AX2 *D. discoideum* cells grown on bacteria was performed according to the same protocol as the fusion of DdB *D. discoideum* cells described in [12]. An amount of 1-2 $\times 10^8$ cells from an overnight shaking culture was washed and resuspended in Sörensen's buffer to a final concentration of $5 \times 10^7$ cells/ml to initiate development. The cells were then allowed to aggregate in buffer droplets with a volume of 1 ml for 2-3 h. Cell aggregates on top of the droplets were transferred to an electroporation cuvette (4 mm gap, VWR). Three square wave pulses of 1 kV for 70 $\mu$s were applied with 1 s intervals between each pulse using an ECM 2001 Electro Cell Manipulator (Harvard Apparatus, Holliston, MA, USA 01746-1388). Afterwards, the cells were immediately transferred to a tube containing 3 ml of Sörensen's buffer with an additional 2 mM $CaCl_2$ and 2 mM $MgCl_2$. After 1 min, 20-50 $\mu$l of cell suspension from the bottom of the tube was transferred for imaging to a 35-mm glass-bottomed microscopy dish (FluoroDish, World Precision Instruments). Wave formation was most abundant 3-6 h after the initiation of development.

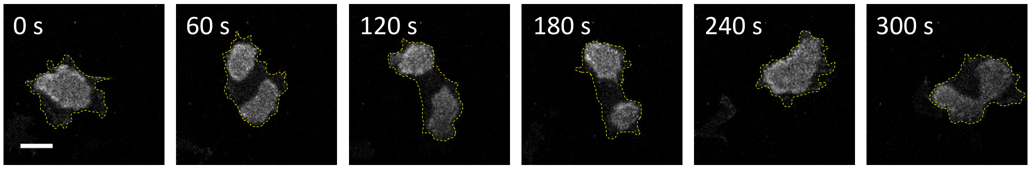


FIG. S5. Meandering and splitting actin wave in the ventral cortex of a normal-sized *D. discoideum* cell.

### S6. LIVE CELL IMAGING

An LSM 780 (Zeiss, Jena) was used for live cell imaging. Excitation was performed with a 488-nm laser (GFP) and a 561-nm laser (RFP). Either a 63× (alpha Plan-Apochromat, NA 1.46, Oil Korr M27, Zeiss, Germany) or a 40× (Plan-Apochromat, NA 1.4 Oil DIC M27, Zeiss, Germany) objective lens was used. Images of the actin cortex were taken in a z-plane close to the surface-attached bottom membrane of the cell, where actin wave dynamics can be observed.

### S7. ACTIN WAVES IN NORMAL-SIZED CELLS

When grown on bacteria, axenic *D. discoideum* cells display the typical macropinocytic actin waves with their characteristic ring-shaped structure and meandering dynamics; see Fig. S5 and SM Movie 1.

---
